# A novel unbiased Linkage Disequilibrium estimator through Random Probing

**DOI:** 10.64898/2026.09.09.750508

**Authors:** Tin-Yu J. Hui, Austin Burt

**Affiliations:** Department of Life Sciences, Silwood Park Campus, Imperial College London, Ascot, UK

## Abstract

The linkage disequilibrium (LD) or correlation of alleles at different loci is a fundamental statistic in population genetics. This work presents a new estimator for the standardised LD measure *r*^2^ between a pair of loci. The Random Probe (RP) estimator involves generating a set of random dummy loci, before calculating the *r*^2^ between each synthetic locus and the two focal loci using one of the existing estimators (even it is known to be biased). The correlation between the two vectors of *r*^2^ is the LD estimate. Computer simulations show promising results, with the RP estimator at least as unbiased as the current Ragsdale and Gravel estimator in most common scenarios. In the more challenging scenarios with skewed allele frequencies and small sample size, RP is preferred by having notably reduced bias, and that the bias is less sensitive to sample size and underlying LD, while maintaining mean squared error comparable to existing methods. The new estimator will most benefit applications relying on accurate measures of LD. Beyond LD, this study stimulates further discussions on whether RP estimator can be generalised to other genetic summary statistics or measures of relatedness.

## Introduction

Research concerning Linkage Disequilibrium (LD) is usually classified into one of the two following categories: those aiming to establish the expected LD induced by evolutionary forces, and those trying to find a good LD estimator from genetic datasets (Sved and Feldman, 1973; Weir, 1979; Hill, 1981; Hayes et al., 2003; Waples, 2006). LD measures the association between alleles at two different loci. In a two-locus two-allele system, let *A* and *a* be the alleles on the first locus, and *B* and *b* on the second, yielding four haplotype combinations. *D* = *p*_*AB*_ − *p*_*A*._*p*_.*B*_ = *p*_*AB*_*p*_*ab*_ − *p*_*Ab*_*p*_*aB*_ quantifies the deviation of haplotype frequency of *AB* from the product of their respective marginal allele frequences, and thus is a measure of LD. If *D* = 0 then the pair is in Linkage Equilibrium (LE). The range of *D* is severely restricted by the marginal allele frequencies, and thus the standardised measure is generally preferred:

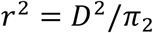

where *π*_2_ = *p*_*A*._(1 − *p*_*A*._)*p*_.*B*_(1 − *p*_.*B*_) is the normalising factor, ensuring *r*^2^ is bounded between 0 and 1 (Hill, 1981).

Thus far we describe LD as observations, or measures of association. In a genomic dataset, we will almost certainly find loci in various degrees of linkage from LE to complete LD. The next step is to associate the observed LD patterns to the evolutionary forces that shape them. Recombination breaks down physical linkage resulting in *E*[*r*] → 0 at a rate of (1 − *c*) per generation (Hill and Robertson, 1968), where *c* is the recombination rate between two loci, also a measure of genetic distance. With finite *N*_*e*_ genetic drift induces correlation by chance such that *Var*(*r*) = *E*[*r*^2^] > 0. For example, 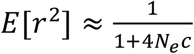 at drift-recombination equilibrium for small *c* (Sved, 1971). Furthermore, LD can be regarded as a probability measure of linked-identical-by-descent (LIBD) (Sved and Feldman, 1973), or covariance of coalescent times between two loci (McVean, 2002). Other processes, such as linked selection, influence the LD pattern between loci under selection and those in the neighbourhood. Hitchhiking occurs when the whole haplotype is dragged along by the increase in frequency of a new favourable mutation (Maynard Smith and Haigh, 1974). Balancing selection maintains long-term LD when the favoured allele combinations are non-randomly associated (Negm and Veller, 2026). Admixture also leaves a footprint on LD (Sved, 2013). Some features shaping chromosome architecture, such as inversions and speciation islands, may suppress local recombination and thus blocks of loci in strong LD are anticipated (Love et al., 2016).

Inference of LD patterns and demographic histories relies on accurate measures of *r*^2^ from genomic datasets. This leads to the second half of the question on the estimation of *r*^2^ from samples. There are numerous fundamental challenges in finding LD estimators. First, with *r*^2^ being a ratio, its unbiased estimators may not be found, even if those for the numerator and denominator exist. Another issue is the systematic bias induced by finite sample size *s*. Third, different estimation routines are required depending on whether gametic phase is known. For phased data, the naïve estimate of *r*^2^ can be calculated directly from the observed haplotype frequencies, which is also the maximum likelihood estimate (mle) under multinomial sampling (Weir, 1979). Unfortunately, it is biased under finite *s*. Another way is to find the unbiased estimators for *D*^2^ and *π*_2_ then find the ratio of the two (Ragsdale and Gravel, 2020, more below). Unphased diploid genotypic data brings a further complication in which the two configurations of the double heterozygote are indistinguishable, resulting in the haplotype frequencies not being directly observable. The question now becomes to estimate haplotype frequencies from genotype counts. With the additional assumptions of random mating and Hardy-Weinberg Equilibrium (HWE), numerical methods and transformations were subsequently developed to solve for mle, although not all mathematical solutions are biologically feasible (Excoffier and Slatkin, 1995; Gaunt et al., 2007; Hui and Burt, 2020). There are also the popular gene-counting Burrows’ method which tolerates some departure from HWE (Weir, 1979), and Rogers and Huff’s (2009) method which relaxes the assumption of random mating. Note that these estimators are known to be biased, or only asymptotically unbiased when *s*−> ∞. It is suggested that the bias goes like 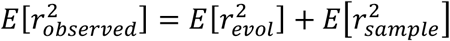, implying the observed *r*^2^ can be partitioned into two additive components: that induced by evolutionary forces and that by finite sampling (Waples, 2006), from which the former part is of our main interest. It is generally understood that bias due to sampling is in the order of 1/*s* (Hill, 1981; Waples, 2005), which easily dominates the observed signal when *s* is small. More recently, Ragsdale and Gravel (2020) expressed the unbiased estimators for *D*^2^ and *π*_2_ as polynomials of the nine genotype counts (∼200 terms in total), and sensibly 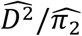 is an estimator for *r*^2^ (the RG estimator). Although 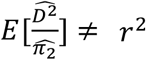, it was demonstrated to be less sensitive to *s* compared to other methods.

## Methods

We propose a new *r*^2^ estimator between a pair of biallelic loci *A* and *B*, with the objectives being it is reasonably unbiased and precise (low variance). The procedure is surprisingly intuitive:

1. Beyond loci *A* and *B*, a set of *L* random probe (or dummy) loci {*Z*_1_, *Z*_2_, …, *Z*_*L*_} needs to be generated. For phased haplotypes, *Z*_*i*_ is formed by random sampling of the “0” and “1” alleles, or by further reshuffling. For unphased diploids, two random alleles are added together to form a genotype such that HWE is largely followed.
2. Probing. For each *Z*_*i*_, find its *r*^2^ between loci *A* and *B* respectively, using an existing estimator of your choice (e.g. Burrows, RG), even if it is known to be biased. You will have obtained a pair of 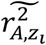 and 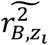.
3. After performing the same for all *L* loci, find the sample Pearson correlation between the two vectors 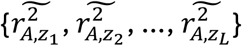 and 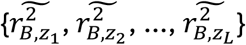, and this is an estimate for 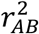. We call this the Random Probe (RP) estimator.

Note that the above procedure applies to both phased and unphased data. Extensive simulations were run to examine the properties of RP under various combinations of true marginal allele frequencies and *D*, as inspired by Ragsdale and Gravel (2020). In each scenario, repeated sampling was conducted, and *r*^2^ was re-estimated from the chosen estimators. Mean square error (MSE), bias, and variance were used to evaluate the estimators as a function of sample size. Another objective is to compare RP against the leading “unbiased” RG estimator. A second set of simulations was run to find out how the number of dummy loci *L* affect precision.

## Results

The performance of RP is summarised in Figures 1 and 2. Three scenarios, representing loci in LE, stronger LD, and skewed marginal allele frequencies, were examined. For phased data (Figure 1), the naïve MLE was also included alongside RG and RP. MLE was severely biased upward under small *s* across all three scenarios. RG and RP performed almost identically in the first two scenarios. In the third scenario with skewed allele frequency at the one locus, MLE significantly overestimated *r*^2^ while SL underestimated it. The two versions of SL lied between the two, with bias about half of RG’s at small *s* ≈ 50. MLE usually had the lowest variance, followed by RG. Note that RP’s variance gradually converged to MLE’s as *s* decreased. Again, one exception was with skewed allele frequencies, where the RP variance was a fixed ratio above that of MLE. All estimators shared similar MSE, a measure of total error.

**Figure 1.**
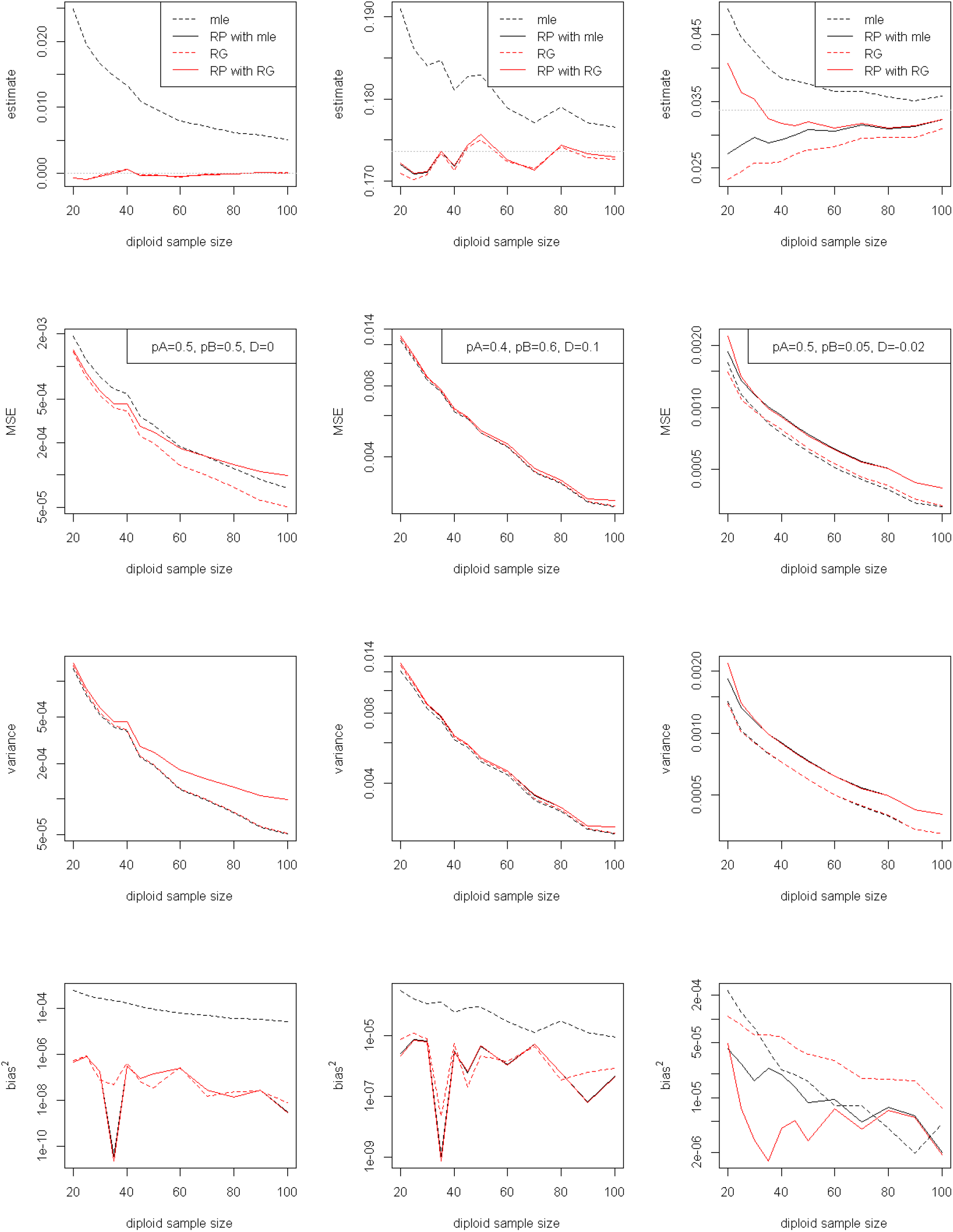
Simulation results for phased data as a function of diploid sample size *s*, ranging from 20 to 100. The three columns represent the three scenarios considered (see true marginal allele frequencies and *D* on the second row). Four metrics were computed to assess the performance of the estimators: (top to bottom) *r*^2^ point estimate, with true value shown in grey, MSE, variance, and bias^2. Under each *s*, 2000 independent repeats were run, and the average was reported. *L* = 20,000 was used. MLE and RG were included for comparison. Note that in some cases the two versions of RP performed almost identically hence often they are visually indistinguishable.

**Figure 2.**
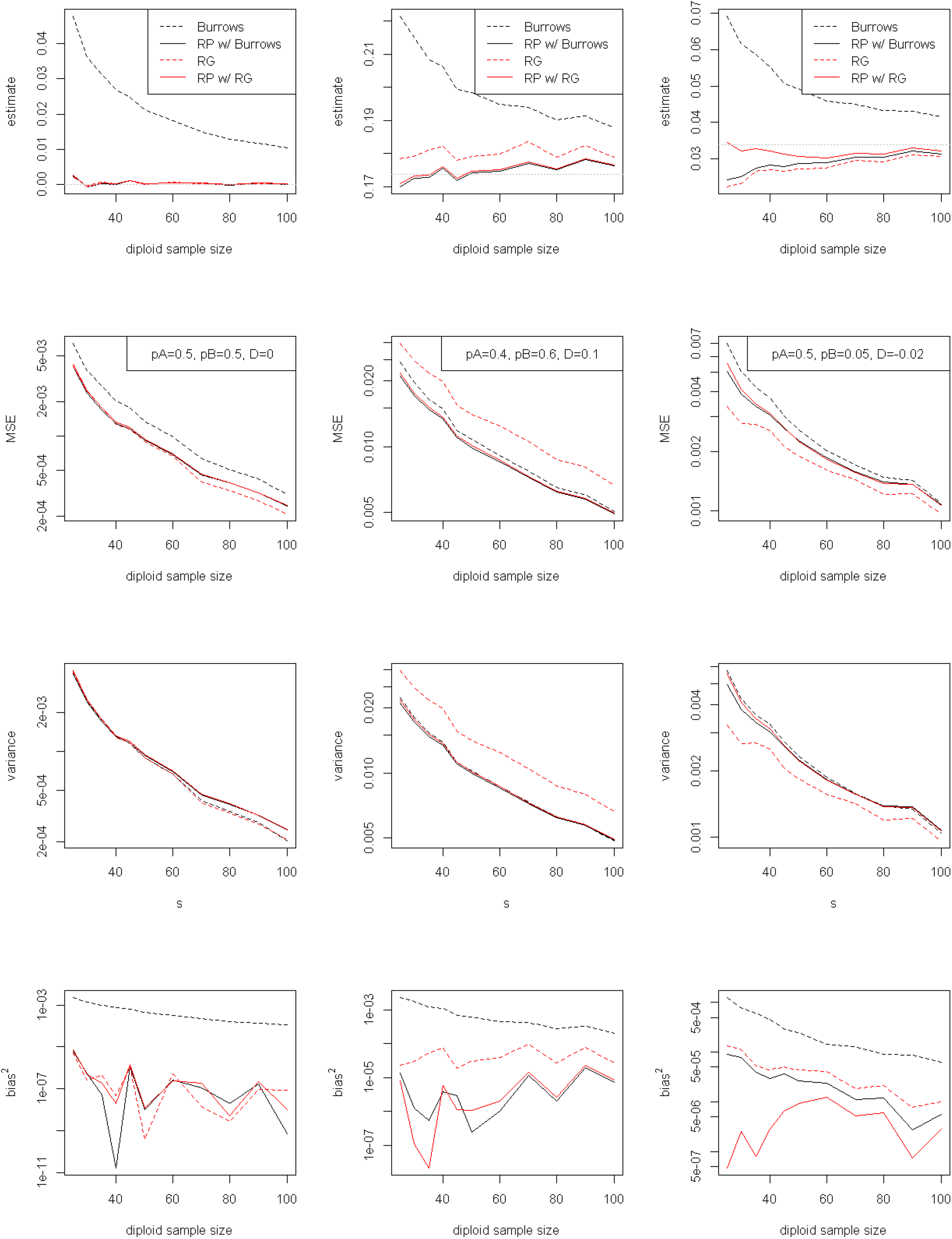
Simulation results from unphased data as a function of s. Burrows’ *r*^2^ and RG were included for comparison. Other settings were the same as Figure 1. The RG estimator was implemented by converting the Mathematica notebook from the supplementary information of Ragsdale and Gravel (2020).

Unphased data gave very similar patterns in that RP was at least as unbiased as RG (Figure 2). Burrows’ *r*^2^ was also included for comparison, and it consistently overestimated *r*^2^, with bias remaining substantial even for large *s* ≥ 100. Burrows’ bias^2 was at least an order magnitude larger than the other two. RG and RP were notably less sensitive to *s*, with RP slightly better across the three scenarios. RP once again outperformed RG under skewed allele frequencies (Figure 2, third column) from which RG tended to underestimate *r*^2^ for small *s*. The variances and MSE of the estimators varied across scenarios: Under LE (Figure 2, first column), RG and RP were equally efficient achieving the lowest variance. For skewed allele frequencies, RG had the lowest MSE despite not having the smallest bias. There exist scenarios (Figure 2, second column) that RP had the lowest variance and bias. RP relies on an existing (biased) estimator to compute LD with the dummy loci. While no specific choice was required, it appeared that pairing the already less-biased RG with RP gave the most consistent results to further remove residual bias.

In the second set of simulations, we examined the precision (1/variance) of the RP estimator as a function of the number of dummy loci *L*. Precision increased with *L*, before reaching a plateau at *L* > 5000 (Figure 3). In addition, *L* only affected the precision but not the bias of RP. The results were almost identical for phased and unphased data.

**Figure 3.**
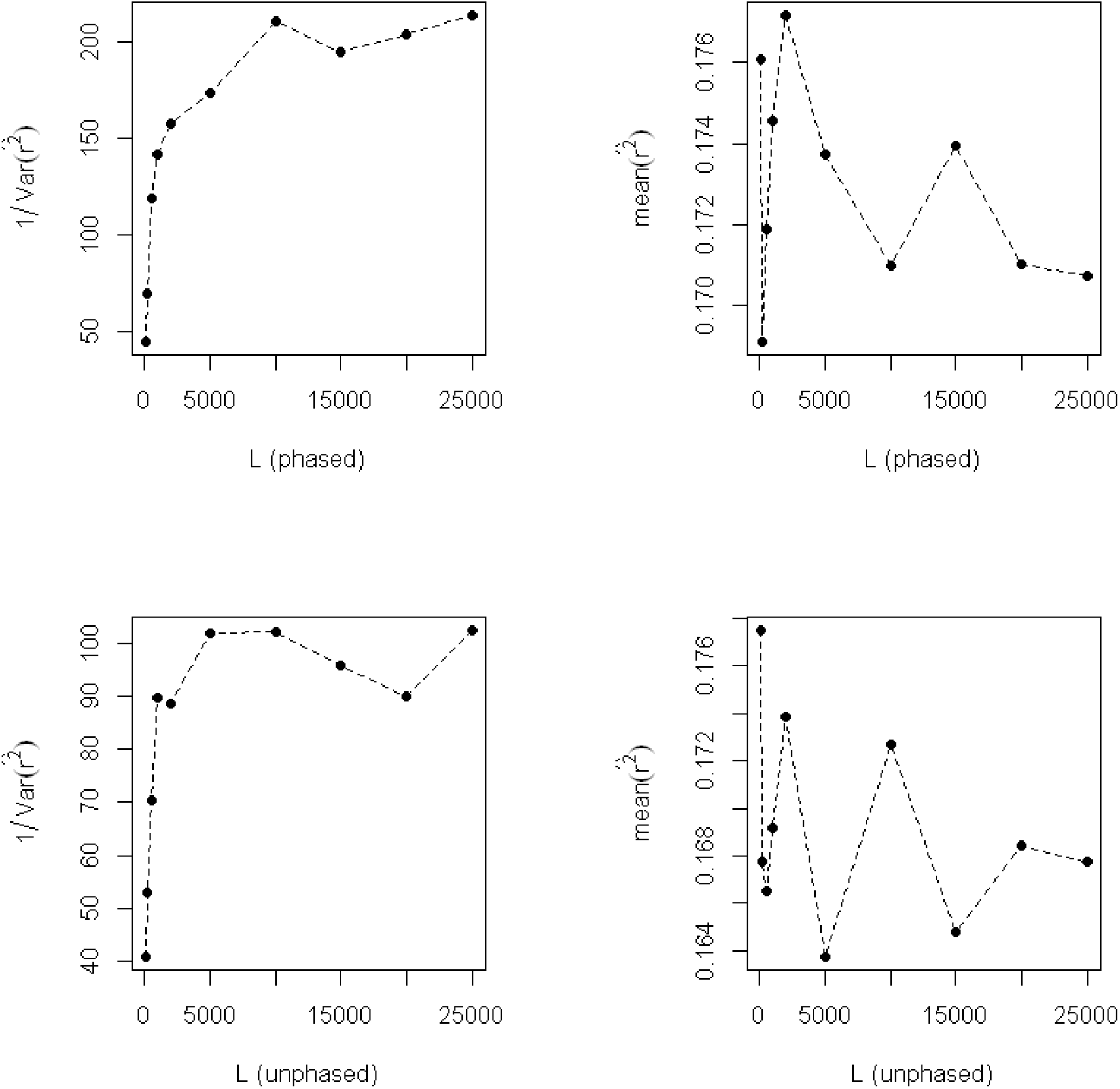
Precision (1/variance, left column) and mean of *r*^2^ (right column) of RP as a function of *L*. Under each *L*, 1000 independent repeats were simulated. Both phased (top row) and unphased (bottom row) data were examined. The underlying population was assumed to be *p*_*A*_ = 0.4, *p*_*B*_ = 0.6, *D* = 0.1 (see middle column of Figure 1), with *s* = 50.

## Discussion

The evolutionary importance of *r*^2^ is indisputable yet finding good estimators for it remains an on-going challenge. Unbiased estimators for ratios may not exist, or at best can only be approximated (e.g. via the delta method (Gerard, 2021)), even those for the numerator and denominator are obtainable. Unlike all other traditional LD estimators, which reasonably (and rightly) focused on the pair of loci of interest, the RP estimator considers an additional set of randomly generated probe loci and their interactions with the pair. The implementation is straight forward that it reuses an existing estimator even if it is known to be biased, then computes the correlation on these *r*^2^ values. Promising results were obtained, that the bias of RP was minimal throughout, and was amongst the least sensitive to sample size *s*, marginal allele frequencies, and the underlying LD. We then compared it to the leading RG estimator, which has previously been demonstrated to outperform most other existing methods, including EM (to solve mle numerically), in both phased and unphased scenarios (Ragsdale and Gravel, 2020). We found that RP was at least as unbiased as RG when the underlying *D* ≈ 0 and the loci were highly polymorphic. In the more challenging scenarios with skewed allele frequencies (third column of Figures 1 & 2) its ability to further reduce bias makes RP the preferred option. Often the bias was reduced by 50% for small *s*<=50, the parameter range in which bias from sampling is most pronounced. Traditional estimators struggle when the minor allele frequency (maf) and/or *s* are both small, leading to the problematic setting of have very few (or zero) expected counts in some cells of a 3-by-3 genotypic table. RP alleviates the bias by iterating though *L* probe loci such that its estimates do not depend only on a single contingency table but the correlation of multiple tables. Empirical corrections were proposed for Burrows’ *r*^2^ but only for unlinked loci under the context of *N*_*e*_ estimation (Waples, 2006), thus are unable to generalise to broader scenarios. A minimum maf is usually imposed when analysing real datasets (Waples and Do, 2010), however, since our aim was to test the estimators to their limits, all loci were included as long as their observed maf>0. In our experience, the maf of the probe loci did not cause noticeable effect on the bias, nonetheless, we followed the general rule that polymorphic loci are of the most informative, thus they had maf of at least 0.4 in our simulations.

Using too few probe loci will result in large estimating variance (Figure 3). We recommend using *L* > 5000 as the precision is approaching a plateau beyond this value. In theory, for *L* pairs of data points, the bias is of the order of *O*(1/*L*) when estimating the correlation coefficient, but as *L* is set to large (*L* >>> *s*) it becomes negligible. There is some additional computing cost for implementing RP but this can easily be mitigated by efficient programming such as parallelisation. Note that 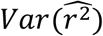 from unphased data is approximately twice that from phased data (Figure 3), in agreement with a previous study (Hui and Burt, 2020). As for most existing estimators, RP assumes multinomial sampling from true (fixed but unobserved) gametic frequencies, and additionally HWE and random pairing of gametes for unphased diploids. Further assumptions may apply to downstream inference of evolutionary processes. This new RP estimator will benefit nearly all applications concerning LD by providing a relatively unbiased measure. Note that an unbiased estimator may not necessarily be the “best”, as in some cases it may be preferable to minimise variance or MSE. In our view, inferences of evolutionary processes will benefit most from having an accurate measure of LD, while the slight increase in estimation variance can be offset by aggregating signals from the tens of thousands of loci found in genomic datasets (e.g. whole-genome sequencing data).

While its empirical performance was thoroughly examined via computer simulations, the rationale behind the method requires further thought. It is obvious that if the two loci are in complete LD, then their *r*^2^ with a third-party should always be perfectly correlated. Beyond measuring association, LD has long been considered as a correlation as well as a probability. Sved and Feldman (1973) proposed the equivalence of *r*^2^ and the probability of LIBD, i.e. the chance of two alleles being inherited together without recombination. If we assume the probe locus are on a different chromosome (from *A* and *B*), then *Z*_*i*_ shares the same LIBD probability with the two. The triplet however is not always inherited together, as this also depends on whether *A* and *B* are in LIBD. If we consider an array of *Z*_*i*_’s, and count the concordance and discordance pairs of whether the three are inherited together as a correlation measure, then it should contain information about the LIBD relationship (thus *r*^2^) of the two focal loci. Another consideration is *Cor*(*a* + *bX, c* + *dY*) = *Cor*(*X, Y*) for random variables *X* and *Y*, and constants *a, b, c, d*. Given the goal of our first argument is to estimate the correlation between two true 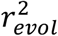, and that the bias due to sampling is approximately linear (Waples, 2006; Hui and Burt, 2020), this explains why RP is relatively unbiased even when the *r*^2^ between the dummy loci are biased. Viewing from another perspective, RP quantifies the correlation between 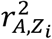 and 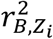 (see step #3 of Methods), which in-turn allows us to account for pseudo-replication when calculating LD from genomic datasets. An in-depth analysis is provided separately (Hui et al., in prep).

It is also worth considering whether the same principle of random probing can be generalised to other population genetic summary statistics, especially those calculated from a pair of individuals, loci, or populations, or for measures of relatedness. Preliminary study shows that probing with a third random population holds information about genetic differentiation (i.e. correlation of allele frequencies) between two focal populations, interesting enough to encourage further investigation.

## Acknowledgements

This work is supported by Wellcome Trust (224487).

## Computer codes

Computer codes will be available soon.

## Competing interests

Authors declare no completing interests.

## Notes

### Competing Interest Statement

The authors have declared no competing interest.

